# Age effects aggressive behavior: RNA-seq analysis in cattle with implications for studying neoteny under domestication

**DOI:** 10.1101/2020.08.22.262493

**Authors:** Paulina G. Eusebi, Natalia Sevane, Thomas O’Rourke, Manuel Pizarro, Cedric Boeckx, Susana Dunner

## Abstract

Aggressiveness is one of the most basic behaviors, characterized by targeted intentional actions oriented to cause harm. The reactive type of aggression is regulated mostly by the brain’s prefrontal cortex; however, the molecular changes underlying aggressiveness in adults have not been fully characterized. Here we used an RNA-seq approach to investigate differential gene expression in the prefrontal cortex of bovines from the aggressive Lidia breed at different age stages: young three-year old and adult four-year-old bulls. A total of 50 up and 193 down-regulated genes in the adult group were identified. Furthermore, a cross-species comparative analysis retrieved 29 genes in common with previous studies on aggressive behaviors, representing an above-chance overlap with the differentially expressed genes in adult bulls.

Particularly, we detected changes in the regulation of networks such as synaptogenesis, involved in maintenance and refinement of synapses, and the glutamate receptor pathway, which acts as excitatory driver in aggressive responses. Our results provide insights into candidate genes and networks involved in the molecular mechanisms leading to the maturation of the brain. The reduced reactive aggression typical of domestication has been proposed to form part of a retention of juvenile traits as adults (neoteny). The significant age-associated differential expression of genes implicated in aggressive behaviors and concomitant increase in Lidia cattle aggression validates this species as a novel model comparator to explore the impact of behavioral neoteny under domestication.

## Introduction

Aggression in animals, including humans, serves important purposes in securing mates, territory and food and, therefore, is one of our most basic behaviors. We understand aggression as an overt behavior directed at an object or a subject with an intention to cause harm or damage (Gannon et al. 2009). This definition encapsulates a heterogeneous and multifaceted construct, including the important distinction between two different subtypes of aggression, reactive and proactive (Smeets et al. 2017). Reactive aggression is the impulsive response to provocations, frustrations or threats from the environment. Proactive aggression, is conscious and goal-directed, planned with a specific outcome in mind (e.g. attainment of resources). While both subtypes can co-occur, different neurobiological structures have been shown to underlie reactive and proactive subtypes (Blair, 2013).

Animal studies have shown that reactive aggression is mediated by the limbic system, through a circuit that runs from the amygdala to the hypothalamus, and from there to the periaqueductal grey. This circuitry is regulated by frontal cortical regions, particularly by the prefrontal cortex (PFC) and the anterior cingulate cortex (Blair, 2013). There is strong evidence that reactive aggression often appears earlier in life than proactive aggression, and that heightened levels of reactive aggression at early stages are associated with increased levels of anxiety in adulthood. In humans, childhood reactive aggression can be found in different neuropsychiatric that can lead to conduct disorders during adulthood (Fite et al., 2010). Therefore, a comparison of gene expression profiles in PFC tissues at different stages of development may be relevant to reveal some of the molecular characteristics and mechanisms involved in aggressive-related perturbations that characterize early stages of aggression and its progression into adulthood.

Research studies across different fields suggest that modern humans have undergone positive selection for reduced reactive aggression relative to our archaic counterparts. Behavioral comparisons with our closest extant ape relatives have shown that instances of reactive aggression in our species are between two and three orders of magnitude lower (Wrangham, 2019). Modern humans also have marked reductions in hypothalamic-pituitary-adrenal (HPA) axis activity (which regulates stress responses) compared to other primates (Chrousos et al., 1982). Furthermore, evidence of convergent phenotypical changes in anatomically modern humans and domesticated species strongly suggest that such reductions in aggressive behaviors have continued in our recent evolutionary history, marking an important contributor to the modern—archaic split: domesticated species ubiquitously display reductions in reactive aggression when compared to their wild counterparts, often accompanied by broad phenotypical changes known as the “domestication syndrome” (reductions in cranial, brain and tooth size, shortening of muzzles, depigmentation, the development of floppy ears, and attenuated HPA activity) (Wilkins et al., 2014). Experimentally controlled selection for reduced reactive aggression in farm-bred foxes brought about many of these broader changes (Dugatkin and Trut, 2017). Modern humans exhibit craniofacial changes relative to Neanderthals reminiscent of those in domesticated species, including reductions in tooth size, contraction of the skull (in our case giving a more globularized shape), and flattening of the face (Sánchez-Villagra & van Schaik, 2019; Theofanopoulou et al., 2017). Genes identified within selective-sweep regions in modern humans have been shown to have above-chance intersection with those implicated in domesticate evolution, including disproportionate targeting glutamate receptor genes associated with aggressive behaviors and implicated in regulation of the HPA axis (O’Rourke & Boeckx, 2020; Theofanopoulou et al., 2017). This evidence for (self-) domestication in ours and domesticated species has long been considered an instance of retained juvenile (neotenic) traits (see e.g. 5). This includes behavioral traits such as diminished reactive aggression. A corollary of this proposal is that Lidia cattle, for which reactive aggression increases with maturity, should have non-juvenile gene expression patterns.

The Lidia cattle breed is a promising and natural animal model for studying reactive aggression, given that the breed’s agonistic responses have been exacerbated through long-term selection of aggressive behaviors. In relation to other domesticated cattle breeds, the selection regime that Lidia cattle have undergone is analogous to that practiced in the farm-fox experiment, where distinct fox lineages have been bred for tame versus aggressive behaviors. Although not perhaps the most obvious models of human evolution, genes targeted by selective sweeps in tame cattle (like dogs and unlike cats and horses) show above-chance intersection with genes falling within selective sweep regions in modern humans (Theofanopolou et al., 2017). The Lidia is a primitive breed, raised for centuries with the exclusive purpose of taking part in socio-cultural and anachronistic events grouped under the term of “*tauromaquias*” (Eusebi et al., 2018). At these festivities, male bovines are placed on a bullring in solitary with a human subject, resulting in a series of aggressive reactions that measures the animal’s fighting capacities by means of reactive responses, such as fighting, chasing and ramming. Different types of *tauromaquia* take place at the bullrings, perhaps the most popular are those known as major festivities: the “*corrida*” and the “*novillada*” (Domecq, 2009). The difference between these two events is the age of the bovines. In a “*corrida*”, the protagonists are 4-year-old bulls, while a “*novillada*” uses 3-year-old bovines (Maudet, 2010). This age discrimination between events is not random; 3-year-old animals are smaller, less strong and their reactive responses tend to be less explosive and intense than those of 4 year-old bovines.

Our previous studies using Lidia bovines as a model to study the neurobiological basis of aggressive behaviors revealed widespread changes in PFC gene expression compared with tamed Wagyu bovines, a meat-production breed known for their docile-temperament. Thus, we aimed to provide insights into potential molecular and cortical substrates of aggressiveness (Eusebi et al., 2020). Furthermore, genome scans for selection signatures in the Lidia cattle breed have uncovered the existence of several genes strongly associated with aggressive behaviors previously reported in humans and other animal models (Eusebi et al., 2019). Particularly, we have associated a novel polymorphism in the promoter of the monoamone oxidase A (*MAOA*) with different levels of aggressiveness among cattle breeds, a gene that has been clearly linked to aggression in humans (Brunner et al., 1993; Eusebi et al., 2019).

Although studies characterizing gene expression differences between juveniles and adults in the PFC have shown promising results in mice (Agoglia et al., 2017; Lander et al., 2017), to our knowledge, there is a lack of such studies using cattle as a model. Thus, the aim of the present study is to explore differences in gene expression at two age stages of Lidia cattle, “young” three-year-old and “adult” four-year-old bovines, given the marked differences in the intensity of their agonistic responses. For this purpose, gene expression profiles from the PFC tissue of eight adult and eight young Lidia bovines were compared. The results of the present study will give insights into the molecular modulation involved in the complex mechanisms leading to the maturation of the brain, with a focus on aggressive behaviors. From an evolutionary perspective, comparisons identifying developmental changes in gene expression can inform as to the possible influences of transcriptional neoteny on the retention of juvenile behaviors into adulthood (Somel et al., 2009). Domestic animals such as dogs retain playful juvenile behaviors as adults, and the reduction of reactive aggression under domestication may also result from a more juvenile neurotranscriptome. Increased aggression of four-year-old Lidia cattle may be expected to rely on an expression profile involving genes not typically implicated in domestication.

## Material and Methods

### Ethics statement

No special permits were required to conduct the research. All animals were sacrificed for reasons other than their participation in this study. Harvesting of bovines’ brains at the cutting room followed standard procedures approved by the Spanish legislation applied to abattoirs (BOE, 1997).

### Animals

Post-mortem PFC tissue samples were retrieved from 16 non-castrated bovine males of the Lidia breed, eight three-year-old and eight four-year-old, for the young and adult groups respectively. In standard RNA-seq studies, this level of replication is 2.6-fold higher than the minimum required (3 individuals/group). Similarly to a classic resident-intruder laboratory test, where an unfamiliar animal (intruder) is placed in the territory of another animal (resident) resulting frequently in an aggressive conflict (Koolhaas et al., 2013), each bovine was placed in a bullring and incited to develop reactive aggressive responses for approximately thirty minutes prior to its sacrifice, to measure their fighting abilities (Domecq, 2009). Samples of PFC were collected less than an hour post-mortem and immediately submerged in RNA-later™ (Thermo Fisher Scientific, Madrid, Spain), followed by 24 hours’ storage at 5°C and long-term conservation at - 80°C.

### RNA isolation and sequencing

Total RNA was extracted from PFC samples using the RNeasy Lipid Tissue Mini Kit (QIAGEN, Madrid, Spain), and total RNA concentration and purity were assessed with a Nanodrop ND-1000 spectophotometer (Thermo Fisher Scientific, Spain). To guarantee their preservation, RNA samples were treated with RNAstable (Sigma-Aldrich, Spain), and shipped at ambient temperature to DNA-link Inc. sequencing laboratory (Seoul, Korea) to perform high throughput sequencing. For quality check, the OD 260/280 ratio was determined to be between 1.87 and 2.0. All these procedures were conducted according to the respective manufacturers’ protocols. Individual libraries for each of the analyzed bovines (*N*=16) were processed using the TruSeq Stranded mRNA Preparation kit (Illumina, San Diego, CA, USA) according to manufacturer’s instructions. Each library was pair-end sequenced (2×100 bp). Individual reads were de-multiplexed using the CASAVA pipeline (Illumina v1.8.2), obtaining the FASTQ files used for downstream bioinformatics analysis. Illumina reads generated from all samples have been deposited in the NCBI GEO / bioproject browser database (Accession Number: GSE149676).

### Bioinformatics analyses

Quality control and pre-processing of genomic datasets was carried out with the PRINSEQ v. 0.20.4 software (Schmeider and Edwards, 2011). To improve the downstream analysis, low quality (Q < 20) and ambiguous bases (N) were first trimmed from both ends of the reads and, then, the trimmed reads were filtered by a Phred quality score (Q ≥ 20 for all bases) and read length of ≥ 68 bp. The obtained FASTQ files were mapped to the bovine reference genome (Bos taurus ARS.UCD 1.2) with the STAR Alignment v.2.7.3a software (Dobin et al., 2013), using default parameters for pair-end reads and including the Ensembl *Bos taurus* ARS-UCD 1.2 as reference annotation. Once reads were mapped, the expression of each mRNA and their relative abundance in fragments per kilobase of exon per million fragments mapped (FKPM) were measured with Cufflinks v.2.2.1 (Dobin et al., 2013). Only expressed transcripts with FKPM values <0 were used for downstream gene expression analyses, leaving a total of 14 animals (eight young and six adults).

The analysis of differential gene expression between young and adult groups was performed using Cufflinks software (Trapnell et al., 2010). The assembled transcripts from all samples were merged using the Cuffmerge command. Cuffdiff was then used applying default parameters and the option “fr-firststrand” to define pair-end reads. A Benjamini-Hochberg False Discovery Rate (FDR), which defines the significance of the Cuffdiff output, was set as threshold for statistically significant values of the Differentially Expressed Genes (DEG). The results of the Cufdiff RNA-seq analysis were visualized with the R software application CummeRbund v.2.28.0 (Goff et al., 2012).

To study the relationship between differences in PFC gene expression among groups and its biological function, we separated the results of DEG in two independent gene-lists according to their Log_2_ Fold Change (FC) in the adult group: up-regulated for those transcripts displaying a Log_2_FC ≥ 0.1; and down-regulated for those with a Log_2_FC ≤ 0.1.

### Cross Species Comparative Analysis (CSCA)

We conducted a comparison between our DEG and a compendium of genes associated with aggressiveness and previously identified in different genomic studies in humans, rodents, foxes, dogs and bovines, as proposed by Zhang-James et al. (Zhang-James et al., 2019). This list is based on four categories of genomic evidence: a) genome wide association studies (GWAS) in humans of different age groups, children and adults (Fernández del Catillo and Cormand, 2016); b) genes showing selection signatures previously identified in the Lidia breed (Eusebi et al., 2018;, Eusebi et al., 2019); c) RNA-seq studies in rodents (Clinton et al., 2011;, Malki et al., 2014) and silver foxes (Kukekova et al., 2011, Kukekova et al., 2018); and d) genes identified in studies with causal evidence of the Online Mendelian Inheritance in Man (OMIM) and the Online Mendelian Inheritance in Animals (OMIA) databases, and a report of knockout (KO) studies in mice (Våge et al., 2010; Veroude et al., 2016; Zhang-James et al., 2019). The gene-list and details of the different studies are described in Supplementary Table 1. Bovine official gene names were converted to their human orthologues using biomaRt (Durinck et al., 2005) in order to homogenize our DEG with the cross-species gene list. We assigned a weight value (weighted ranking, WR) with the same conditions proposed by Zhang-James et al. (2019). For statistical analysis of the intersection between DEGs and genes identified in studies of aggression, we cross-referenced each gene list using the PANTHER (www.pantherdb.org), NCBI HomoloGene,(www.ncbi.nlm.nih.gov/homologene) and Ensembl orthologue databases *Bos taurus* ARS-UCD 1.2 and Human reference (GRCh38.p13) genomes. If no human–bovine one-to-one orthologues were found in any database, we removed the relevant genes for statistical analysis.

### Ingenuity pathway analysis

Up and down-regulated DEG lists were imported into the Ingenuity Pathway Analysis (IPA) (QIAGEN, www.qiagen.com/ingenuity) software to assess Gene Ontologies (GOs) and canonical pathways enrichment scores. Additionally, up and down-regulated DEGs in common with the CSCA were imported together to build biological networks and identify upstream regulators.

For the canonical pathway analyses, a right-tailed Fisher’s exact test was performed for the enrichment of the DEGs in IPA’s hand-curated canonical pathway database. Here, the P-value calculated for a pathway measures the probability of being randomly selected from all of the curated pathways. To control the error rate in the analysis, IPA also provide the corrected P-values to identify the most significant results in IPA’s canonical pathways based on the Benjamini-Hochberg method (Benjamini and Hochberg, 1995). This tool allowed us to identify the signaling pathways in which the DEGs were enriched. We used the predetermined cut-off of the corrected P-value < 0.05 to define the significant pathways.

For network enrichment and detection of upstream regulators, the genes in common with the CSCA were overlaid onto a global molecular network (GMN) developed based on the Ingenuity Pathways Knowledge Base, in which functional relationships such as activation, chemical-protein interaction, expression, inhibition and regulation of binding were manually searched. Sub-networks of genes were then extracted from the GMN based on their connectivity using the algorithm developed by IPA. The sub-networks were visualized using IPA’s Path Designer tool.

## Results

### Sequencing and read assembly

For the 16 individually sequenced samples, we generated an average of 61.6 million pairs of 100-bp paired-end reads per sample after filtering by quality score. For each sample, an average of 91.6% reads were uniquely mapped to the bovine reference genome, ranging from 86.8 to 95.2% (Supplementary Table 2). The transcripts expressed in both the young and adult groups reaching FKPM values < 0 (*N*=8 young and *N*=6 adult) were included in the differential expression and subsequent IPA analyses (Supplementary Figure 1).

Gene expression differences revealed a total of 27,153 DEGs between young and adult groups. Of those genes, only 243 were statistically significant; 50 up-regulated (log_2_FC ≥ 0.1) and 193 down-regulated (log_2_FC ≥ −0.1) (Figure 1A and B). For the detailed list of up and down-regulated DEGs see Supplementary Table 3.

**Figure 1.**
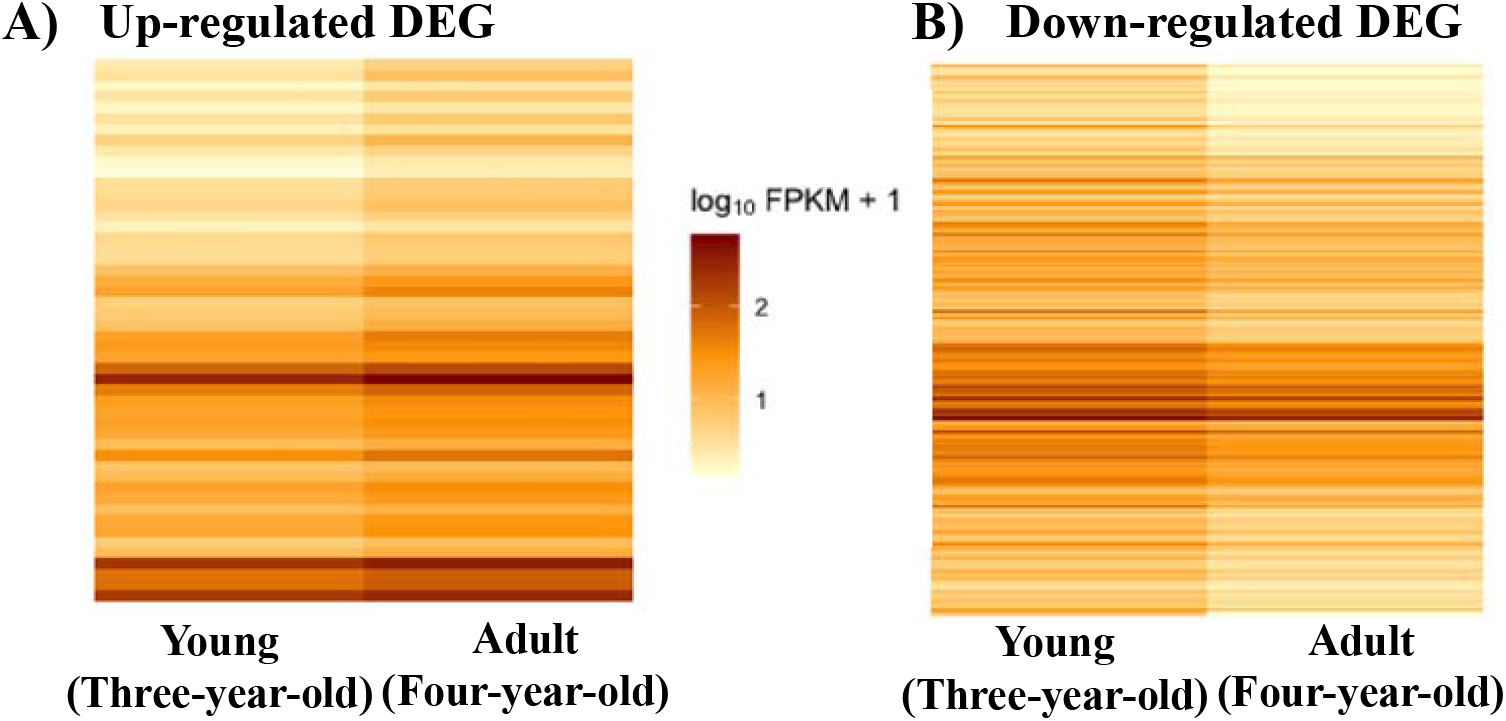
Heatmap of up-regulated (A) and down-regulated (B) differentially expressed genes (DEG) in the adult group.

### Genes in common with the cross-species comparative analysis (CSCA)

The cross-reference analysis between our up and down-regulated DEGs in the adult group and the gene-list associated with aggressive behavior, overlapped at numerous loci. Notably, the down-regulated DEGs shared 21 genes, while the up-regulated displayed only 8 genes in common with the gene reference list (Table 1). The subset of DEGs included in the CSCA are shown in the expression bar plot of Figure 2, which details their standard deviation of FPKM values. As well as these 29 genes, a many-to-many orthologous match was found between the bovine non-classical major histocompatibility complex class I antigen (BOLA-NC1) and the human equivalents (HLA-A, HLA-B, HLA-C, and HLA-F).

**Table 1.**
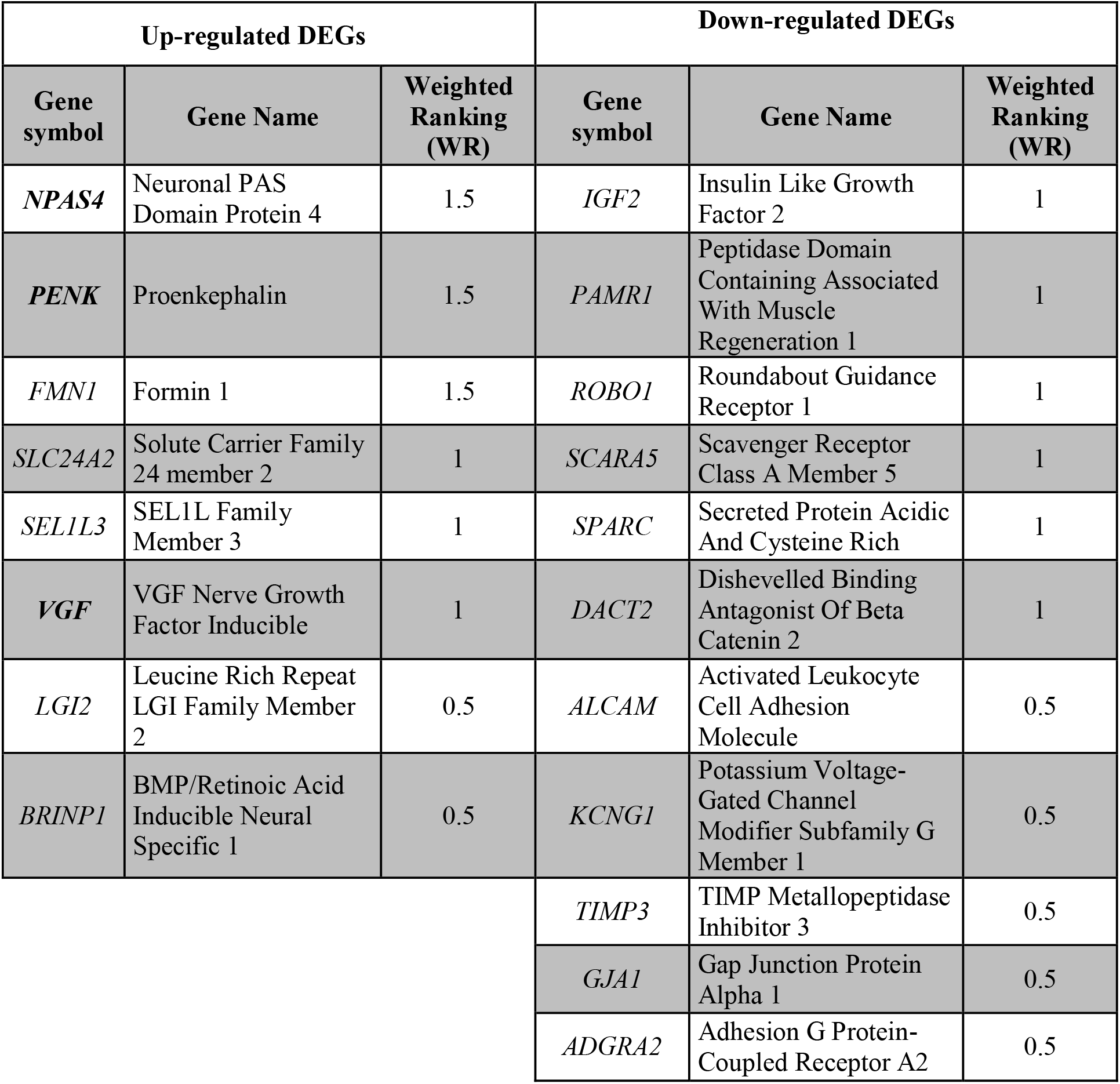
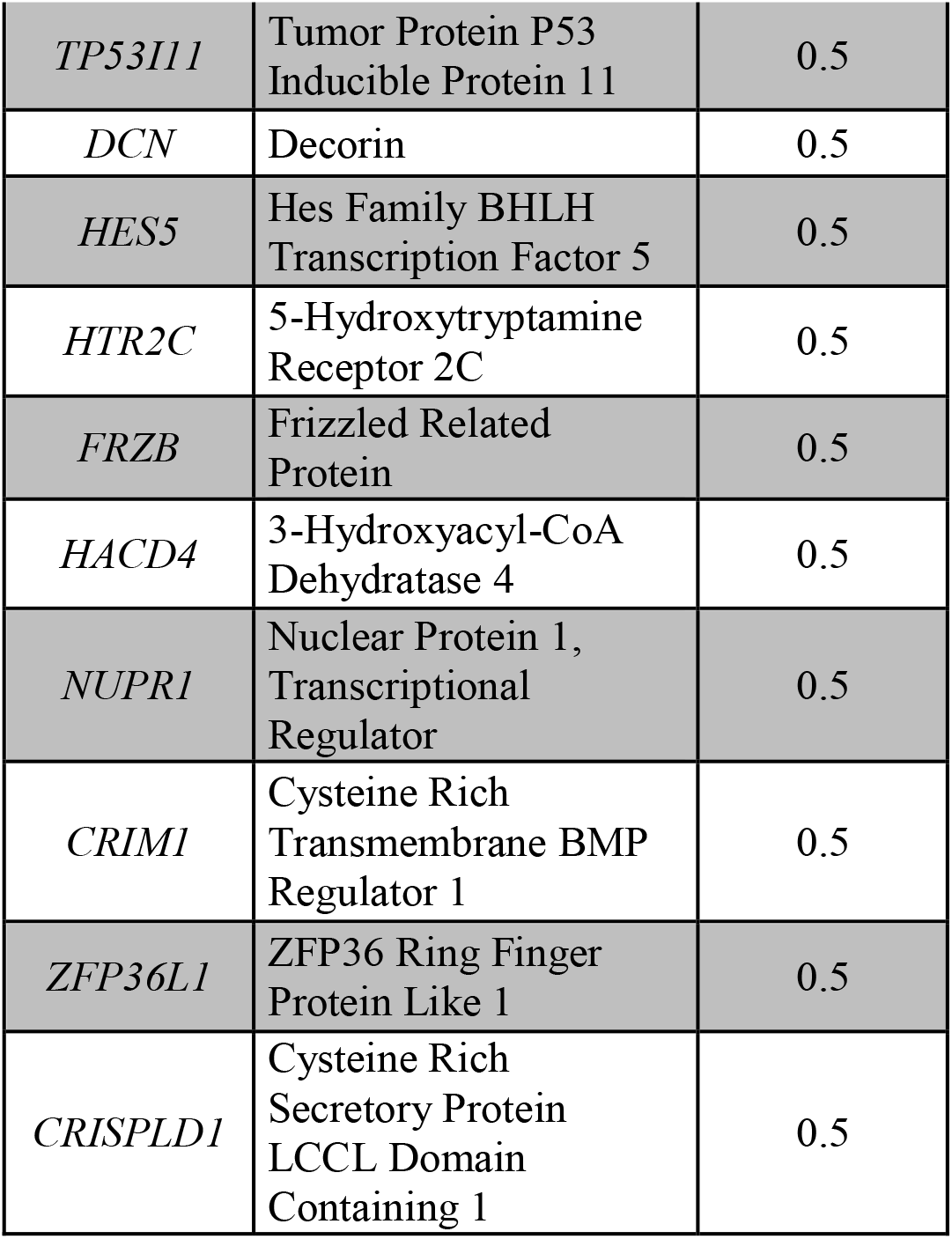
Up and down regulated DEGs in four-year-old bulls in common with the cross-species comparative analysis (CSCA).

| Up-regulated DEGs |  |  | Down-regulated DEGs |  |  |
| --- | --- | --- | --- | --- | --- |
| Gene symbol | Gene Name | Weighted Ranking (WR) | Gene symbol | Gene Name | Weighted Ranking (WR) |
| *NPAS4* | Neuronal PAS Domain Protein 4 | 1.5 | *IGF2* | Insulin Like Growth Factor 2 | 1 |
| *PENK* | Proenkephalin | 1.5 | *PAMR1* | Peptidase Domain Containing Associated With Muscle Regeneration 1 | 1 |
| *FMN1* | Formin 1 | 1.5 | *ROBO1* | Roundabout Guidance Receptor 1 | 1 |
| *SLC24A2* | Solute Carrier Family 24 member 2 | 1 | *SCARA5* | Scavenger Receptor Class A Member 5 | 1 |
| *SEL1L3* | SEL1L Family Member 3 | 1 | *SPARC* | Secreted Protein Acidic And Cysteine Rich | 1 |
| *VGF* | VGF Nerve Growth Factor Inducible | 1 | *DACT2* | Dishevelled Binding Antagonist Of Beta Catenin 2 | 1 |
| *LGI2* | Leucine Rich Repeat LGI Family Member 2 | 0.5 | *ALCAM* | Activated Leukocyte Cell Adhesion Molecule | 0.5 |
| *BRINP1* | BMP/Retinoic Acid Inducible Neural Specific 1 | 0.5 | *KCNG1* | Potassium Voltage-Gated Channel Modifier Subfamily G Member 1 | 0.5 |
|  |  |  | *TIMP3* | TIMP Metallopeptidase Inhibitor 3 | 0.5 |
|  |  |  | *GJA1* | Gap Junction Protein Alpha 1 | 0.5 |
|  |  |  | *ADGRA2* | Adhesion G Protein-Coupled Receptor A2 | 0.5 |

|  |  |  |
| --- | --- | --- |
| *TP53I11* | Tumor Protein P53<br>Inducible Protein 11 | 0.5 |
| *DCN* | Decorin | 0.5 |
| *HES5* | Hes Family BHLH<br>Transcription Factor 5 | 0.5 |
| *HTR2C* | 5-Hydroxytryptamine<br>Receptor 2C | 0.5 |
| *FRZB* | Frizzled Related<br>Protein | 0.5 |
| *HACD4* | 3-Hydroxyacyl-CoA<br>Dehydratase 4 | 0.5 |
| *NUPR1* | Nuclear Protein 1,<br>Transcriptional<br>Regulator | 0.5 |
| *CRIM1* | Cysteine Rich<br>Transmembrane BMP<br>Regulator 1 | 0.5 |
| *ZFP36L1* | ZFP36 Ring Finger<br>Protein Like 1 | 0.5 |
| *CRISPLD1* | Cysteine Rich<br>Secretory Protein<br>LCCL Domain<br>Containing 1 | 0.5 |

**Figure 2.**
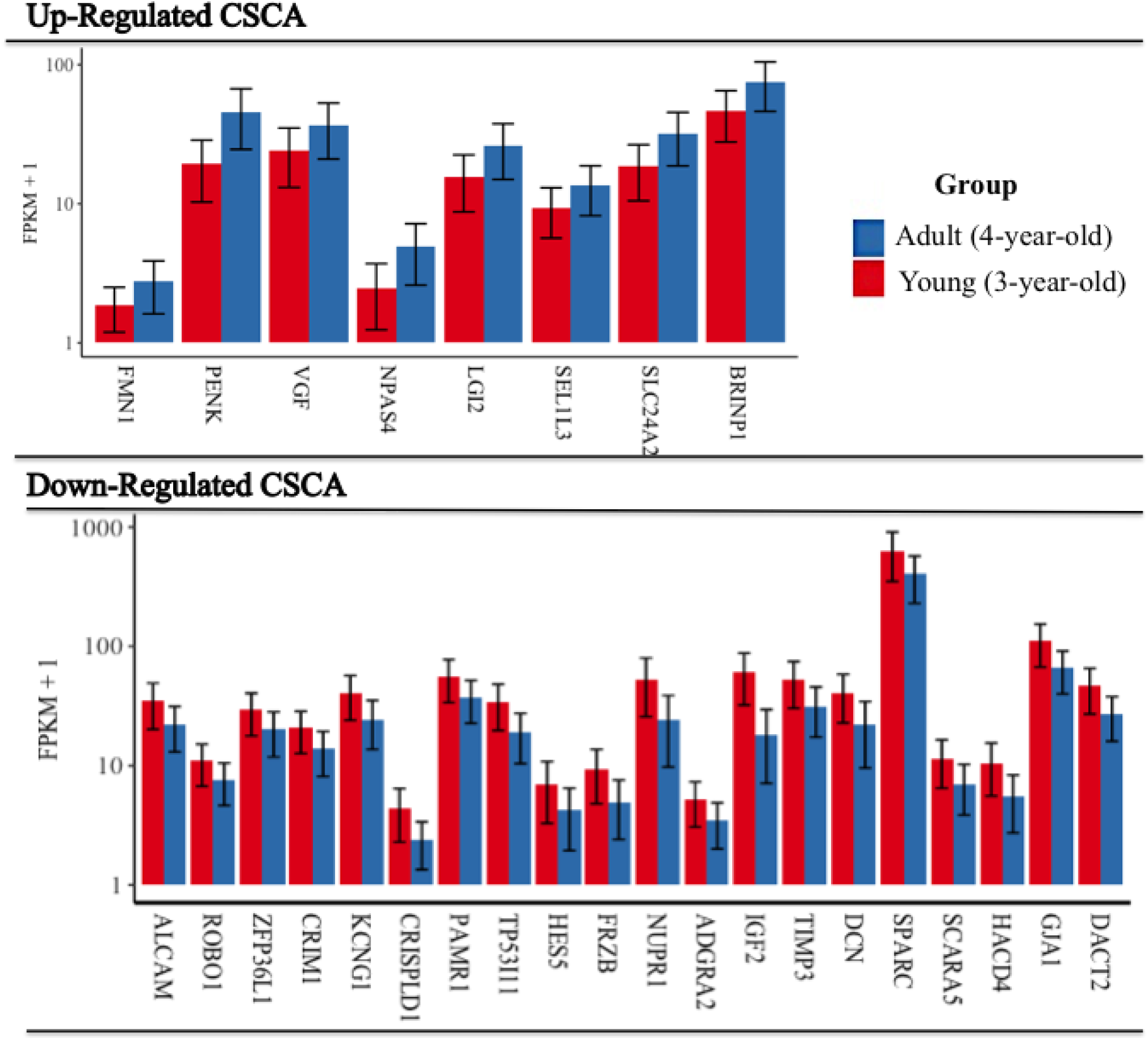
Bar chart of up and down regulated DEGs in common with the cross-species comparative analysis (CSCA). Gene abundance is represented in fragments per kilobase of exon per million fragments mapped (FPKM) of the adult (4-year-old) and young (3-year-old) groups.

### Statistical analysis of aggression-associated differentially expressed genes (DEG)

In order to test whether the 29 adult Lidia DEGs identified in the CSCA represent a statistically significant association between maturation and aggression, we calculated the cumulative hypergeometric probability of this overlap occurring. Following removal of non-orthologous genes, 1701 aggression-associated genes across species and 238 Lidia DEGs remained. Taking the estimated 22,000 genes in the bovine genome (Elsik et al., 2009), an overlap of at least 29 between these two gene samples was significantly unlikely to occur (p-value: 0.0099). We carried out the same analysis taking only aggression-associated genes that were identified in brain-expression studies across the different species (totaling 1151). Of the 238 Lidia DEGs, 21 were also identified in these studies (p-value: 0.0137). Under a more restrictive analysis, whereby only cortically expressed genes are taken as potential comparators ― estimated at 85% of all protein-coding genes in both humans and pigs (Sjöstedt et al., 2020) ―, the intersection of 21 was slightly more likely to have occurred by chance (p-value: 0.0617).

### Ontological analysis of the differentially expressed genes (DEG)

The up and down-regulated DEGs in the 4-year-old adult group were subjected to a Fisher’s exact test within IPA to obtain Ingenuity ontology annotations. IPA annotations follow the GO annotation principle, but are based on a proprietary knowledge base of over a million protein-protein interactions. Significant results are summarized in Supplementary table 4. The most relevant results for the up-regulated DEGs were obtained under the *physiological system development* and the *disease and disorders* categories. Within the first of these categories, the top of the list gathered terms related with *behavior* (highest p-value range of 5.51E-08 and 15 DEG) and *nervous system development and function* (highest p-value range of 7.30E-07 and 24 DEG), while within the category of *diseases*, terms related to *cancer* (highest p-value range of 1.21E-07 and 44 DEGs), *neurological diseases* (highest p-value range of 8.07E-07 and 21 DEGs) and *psychological disorders* (highest p-value range of 3.35E-06 and 12 DEGs) occupied the top positions.

Additionally, we explored the overrepresentation of specific enriched functions within the category of *behavior* and its relationship with diseases or function annotations, finding high overall significance levels of association between up-regulated DEGs and diverse behavioral conditions (p-values < 10^−5^) (Table 2, Figure 3).

**Table 2.** Overrepresentation of enriched specific functions within the category of *behavior*. N: Number of up-regulated DEGs in the adult group.

| Disease / function annotation | P-value | N | Genes |
| --- | --- | --- | --- |
| Emotional behavior | 5.51E-08 | 9 | *BRINP1, CBLN4, GRIN2A, PENK, SCN1B, SLC17A6, SCN4B, SCN1B, SCN1A, VGF* |
| Learning | 3.27E-06 | 9 | *ABCC8, BRINP1, DKK1, GRIN2A, NPAS4, SCN1A, SLC4A2, TAFA2, VGF* |
| Memory | 7.70E-07 | 8 | *ABCC8, BRINP1, DKK1, GRIN2A, NPAS4, SCN1A, SLC4A2, TAFA2* |
| Anxiety | 1.75E-05 | 6 | *BRINP1, CBLN4, GRIN2A, PENK, SCN1A, TAFA2* |
| Spatial learning | 7.09E-05 | 5 | *ABCC8, GRIN2A, SLC24A2, TAFA2, VGF* |
| Irritable behavior | 3.35E-06 | 4 | *GRIN2A, SCN1A, SCN1B, SCN4B* |

**Figure 3.**
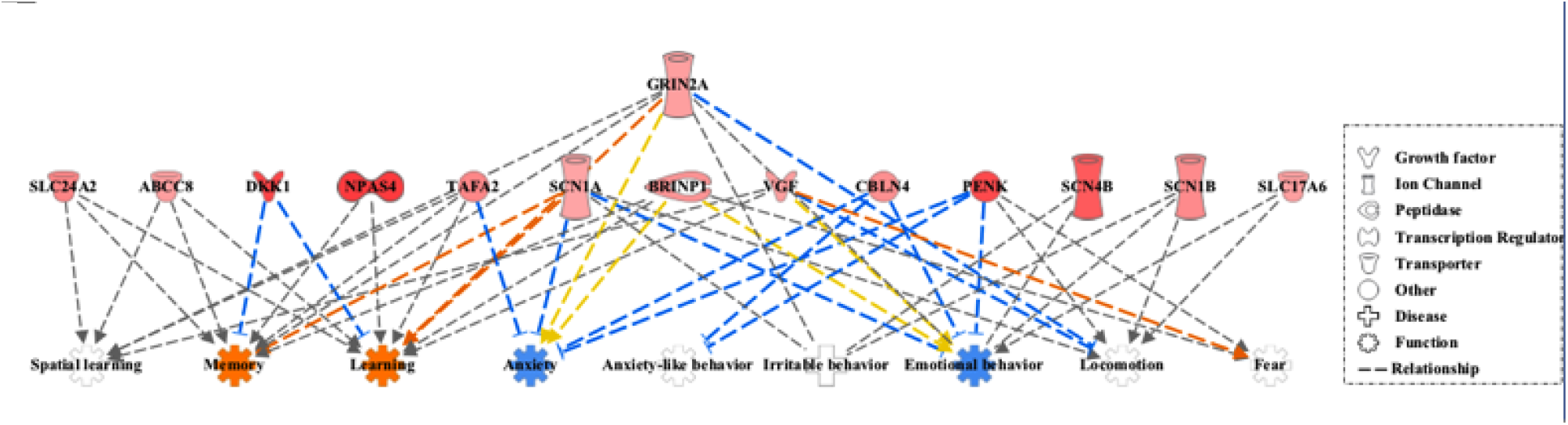
Regulator effects of the IPA package applied to the up-regulated DEG and diverse behavioral conditions (p-values < 10^−5^). In the lower tier, the expected behavioral consequences of the up-regulated DEG changes are shown by considering the Ingenuity Knowledge Base. In orange color are marked those functions predicted to be activated and in blue those predicted to be suppressed.

The Core Analysis of the down-regulated DEGs in the 4-year-old group also revealed genes related to the *physiological system development* and the *disease and disorders* categories. The top enriched genes in the first category were associated with *organismal* (highest p-value range of 2.38E-17 and 63 DEGs), *embryonic* (highest p-value range of 2.38E-17 and 58 DEGs) and *cardiovascular system* (highest p-value range of 1.08E-16 and 43 DEGs) development functions. Furthermore, at the *diseases and disorders* category the down-regulated DEGs were shown to be involved with *cardiovascular disease* (highest p-value range of 3.05E-11 and 34 DEGs), *cancer* (highest p-value range of 1.73E-10 and 84 DEGs), and *organismal injury and abnormalities* (highest p-value range of 1.73E-10 and 84 DEGs) (Supplementary Table 4).

### Metabolic pathways affected by the up and down-regulated DEGs in the adult group

Within the up-regulated DEGs in the adult 4-year-old group, four pathways surpassed the significance threshold (p-values < 0.05, Supplementary Figure 2). The *synaptogenesis* and the *Wnt/*β*-catenin signaling* are among the most significant pathways (p-value = 3.08E-03).

Besides, the *glutamate receptor pathway*, which mediates synaptic signaling by this primary excitatory neurotransmitter in the central nervous system (CNS), resulted also highly significant (p-value = 5.74E-03).

The analysis of down-regulated pathways revealed the involvement of the retrieved DEGs on 53 metabolic routes (Supplementary Table 5). Amongst the most significant, the *intrinsic prothtrombin activation pathway* (p-value =1-08E-08) and, noteworthy, again the *Wnt/*β*-catenin signaling* (p-values = 4.76E-05) route, are highlighted within the down-regulated DEGs in the adult group (Supplementary Figure 1).

### Gene networks and upstream activator analyses of the genes in common with the CSCA

We analyzed the gene transactivation networks and upstream activators of the 29 DEGs with one-to-one orthologous matches in the CSCA, finding three highly interconnected networks related to organ morphology, organismal development, behavior, cancer, neurological disease and psychological disorders (Figure 4).

**Figure 4.**
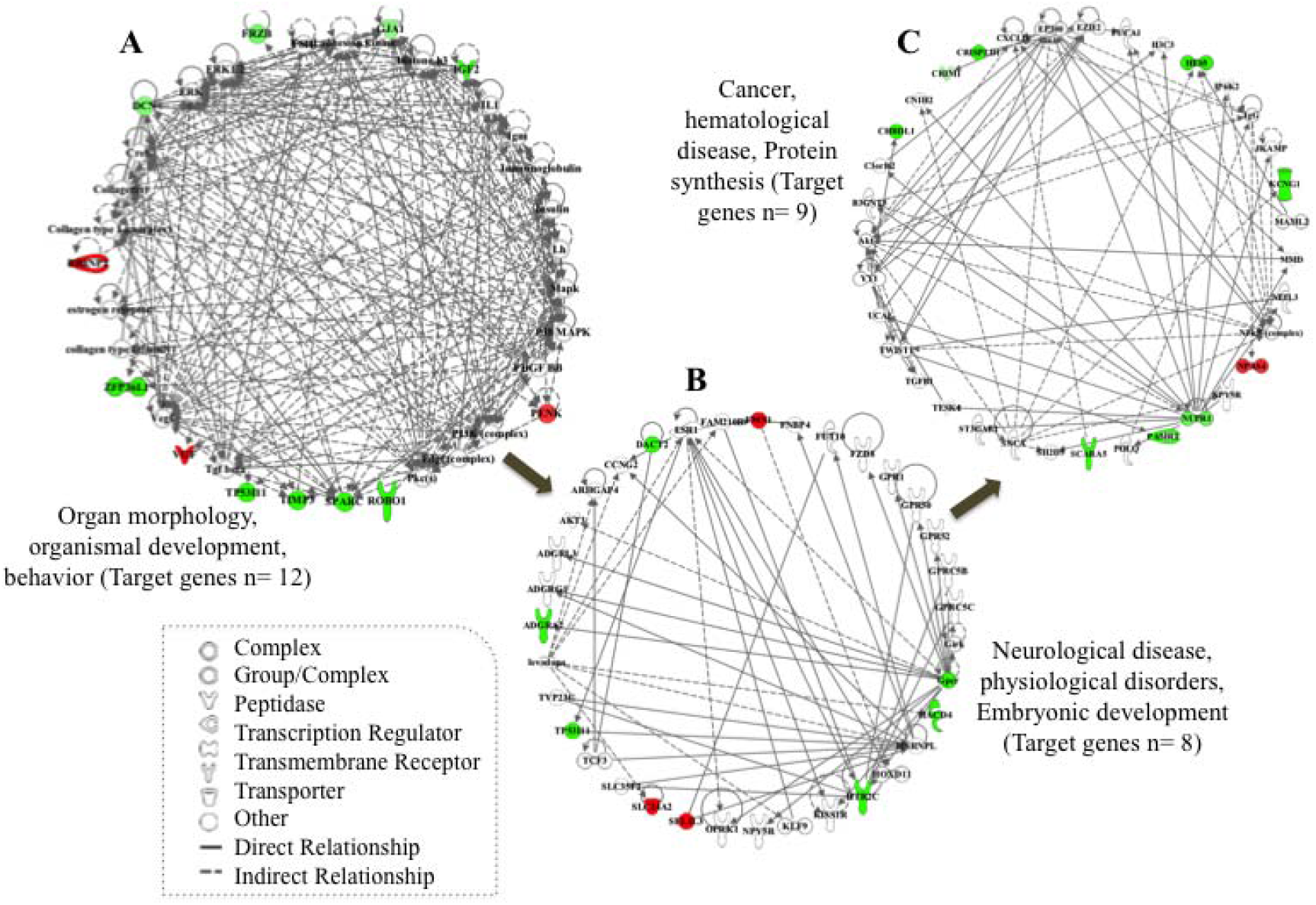
Network analysis of the 29 DEGs in the adult group in common with the CSCA. Genes highlighted in color correspond to the DEGs in our study; in red the up-regulated and in green the down-regulated DEGs.

Finally, the upstream analysis tool of the IPA package was employed to identify potential upstream regulators that may explain the differential patterns of expression between young and adult groups. We explored the activation/inhibition states of the top upstream regulators by using IPA’s activation z-score tool, identifying four main upstream regulators: KRAS proto-oncogene (*KRAS)*, Brain Derived Neurotrophic Factor *(BNDF)*, CAMP Responsive Element Binding Protein 1 *(CREB1)* and SRY-Box Transcription Factor 2 *(SOX2)*. As shown in the graphic network in Figure 5, these genes are involved in an array of diverse biological functions related with behavioral development.

**Figure 5.**
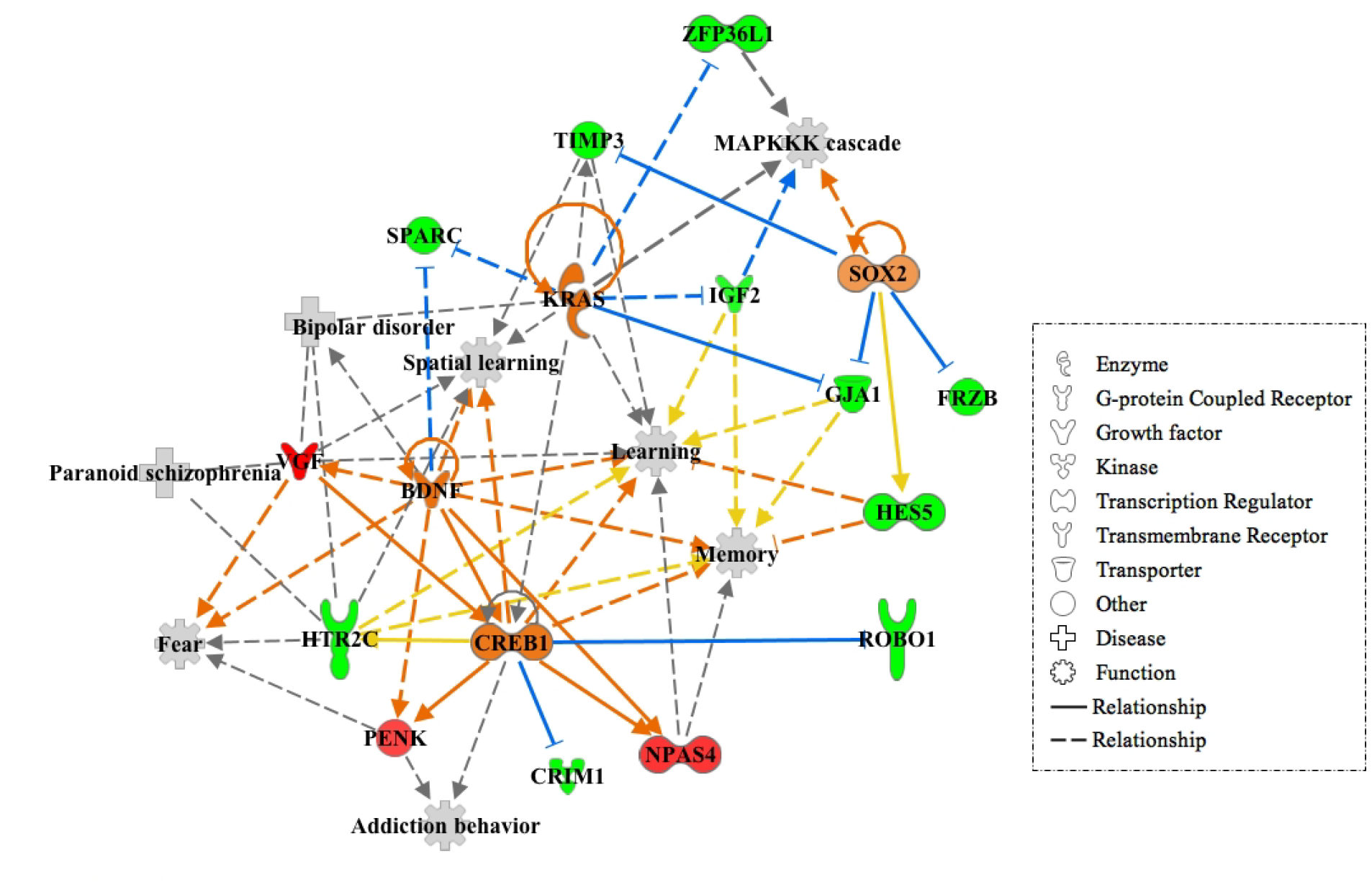
Network visualization of the four major upstream regulators of the DEGs in common with the CSCA. In orange, upstream regulators predicted to be activated; in red and green, genes whose expression increases or decreases in response to the activation of upstream regulation, respectively. The shapes of the nodes reflect the functional class of each gene product, as shown in the legend. The symbols marked in grey define functions. Solid and dashed lines between genes represent known direct and indirect interactions, respectively, with orange lines leading to activation and blue to inhibition.

## Discussion

The structural and molecular changes occurring in the PFC of diverse species at different age stages have been previously explored (Counotte et al., 2010;, Moczulska et al., 2014;, González-Lozano et al., 2016). The majority of these analyses conducted in rodents focus on the study of the molecular changes occurring in post-natal and puberty-adolescence stages.

However, studies on the specific changes of genetic expression during adulthood are still scarce (12, Agoglia et al., 2017, Lander et al., 2017). Here, we used RNA-seq to identify genes and gene networks that show differential expression in the PFC of adult bulls when compared to young bovines of the aggressive Lidia breed.

A total of 143 significant DEGs, 50 up and 193 down-regulated in the adult group were identified. The GO analysis of up-regulated DEGs in the adult group rendered highly significant results for behavioral and nervous system development features, while the down-regulated genes were mostly implicated in systemic developmental functions (organismal, embryonic and cardiovascular system). We detected a modest coincidence between these results and what has been published in previous analysis using high-throughput molecular data in PFC. For instance, Agoglia et al. (2017) used an unbiased proteomics approach to analyze the differential expression of proteins in adolescent compared to adult PFC in mouse. There is a concordance between the proportion of up and down-regulated DEGs in the adult groups reported by both, us and Agoglia et al. (2017), as well as results from the GO analyses showing down-regulation of organismal growth and development processes in adults. This suggests that, both technologies, proteomics and RNA-seq are, as awaited, correlated, detecting similar sets of DE molecules (40). Canonical pathway analyses also revealed that metabolic routes involved in cellular signaling and neurotransmission impact in the adult PFC during the aging process (Figure 2). We detected an over representation of the synaptogenesis and the glutamate signaling reception pathways in the four-year-old bulls, both involved in brain’s synaptic plasticity; the synaptogenesis pathway is implicated in developmental processes of formation, maintenance and refinement of synapses (Cohen-Cory, 2002), while the glutamate signaling receptor acts as excitatory driver of the Hypothalamus-Pituitary-Adrenal (HPA) system, a complex system that triggers reactive aggression responses (Herman et al., 2004; Evanson and Herman, 2015). Similar results were obtained by Lander et al. (2017); they detected an over expression in adults of glutamate markers comparing the cortical mRNA expression of mid-adolescent and adult mice.

Most of the genes making up the glutamate receptor signaling pathway are implicated in NMDA and AMPA-mediated excitation driving long term potentiation at mature synapses. This contrasts with findings that metabotropic and kainate receptor genes ― which tend to down-regulate glutamanergic excitation at synapses, including in limbic circuits that control HPA activity ― are disproportionately implicated in domestication and modern human evolution (O’Rourke & Boeckx, 2020).

In our recent comparison between the Lidia and Wagyu we noted evidence for down-regulated glutamate-to-GABA conversion in the more aggressive breed, suggesting heightened excitatory signaling (Eusebi et al., 2020). In previous studies of selective sweeps distinguishing the Lidia from tamer breeds, we also noted differential signals of selection on the domestication-associated glutamate receptor gene *GRIK3*, further suggesting that heightened excitatory signaling may play a role in the increased reactive aggression of this breed (Eusebi et al., 2018). In future studies it may be worthwhile to compare the age-dependent PFC expression profile of tame cattle breeds with that of the Lidia. Under the hypothesis that increased excitatory signaling contributes to driving aggressive behaviors, one may expect that genes tending to contribute to a relative down-regulation of glutamatergic signaling or delayed maturation of excitatory synapses may be targeted in tamer breeds.

Interestingly, we detected differences in the expression of genes included in the Wnt//β-catenin signaling pathway in both up and down-regulated DEGs in the adult group. The complex interactions of genes, with actors triggering the expression of the Wnt//β-catenin signaling pathway while others are needed to silence routes leading to its suppression, justifies the inclusion of genes from this network in the up and down-regulated cohorts. This pathway is considered one of the most strongly conserved routes among species and is involved in synaptic transmission activities (Maguschak and Ressler, 2012). Malfunction of many components of the Wnt//β-catenin signaling pathway in the adult brain can result in altered behavior and cognitive disorders (Svenningsson et al., 2002).

In addition, the prothrombrin activation pathway acts as signaling route at the CNS, inducing the decrease of cyclic AMP (CAMP) and the activation of protein kinase C (PKC), both enzymes linked to catecholamine and serotonin receptors (David et al., 2004). It has been proposed that, during encounters between individuals, a de-synchronization of the top-down control at these receptors influence a lack of control of emotional reactivity (Kretshcmer et al., 2014). As suggested by Agoglia et al. (2017), the increased expression on these routes observed in the 4-year-old PFC may contribute to the fine-tuning of synaptic transmission, resulting in a superior control of the PFC executive functioning in mature brains, intensifying the addressed aggressive behaviors.

The significant overlap between the age-associated DEGs we identified and the genes associated with aggression in other species strongly suggests an important influence of maturation on the development of aggressive traits in Lidia cattle. This contrasts with the retention of a docile temperament from adolescence to adulthood in other cattle breeds and marks the Lidia as a viable model for research into the neurobiological basis of behavioral neoteny under domestication. Complementarily to these results, the IPA network tool applied to the 29 DEGs with one-to-one orthologues in the CSCA, retrieved three inter-connected networks associated with organismal morphology and development, behavior, cancer, neurological disease and psychological disorders (Figure 3). Particularly, the Mitogen Activated Protein Kinase (*MAPK*) and two Extracellular Signal Regulated Kinases (*ER*K) appear to be involved in diverse molecular mechanisms underlying aggression. Similar results were obtained in previous studies on PFC gene expression (Zhang-James et al., 2019; Eusebi et al., 2020), identifying a differential expression of the ERK/MAPK signaling pathway in their comparisons between aggressive and non-aggressive individuals.

At the regulatory level, three of the four genes predicted to be activated display clear roles in aggressive behaviors, including *BDNF* (Kretshcmer et ak., 2014;,Vigers et al., 2012), *CREB* (David et al., 2004) and *SOX2* (Gatewood et al., 2006). The behavioral functions affected by these regulators include learning and memory cognitive processes, along with behavioral conditions and neuropsychiatric disorders, such as fear, addictive behaviors, schizophrenia and bipolar disorder, all displaying aggressiveness (Figure 4). In this regard, there is evidence of increased memory encoding during early adulthood (Rubin et al., 1998) and also, it has been observed that adults maintain elevated plasma levels of molecules associated with abnormal aggressive behaviors (i.e. corticosterone) longer than adolescents (Romeo et al., 2006).

Furthermore, some neural receptors at adult PFC have shown heightened expression levels, conferring them higher risk of addictive and impulsive behaviors (Sonntag et al., 2014). This evidence suggest that behavioral and physiological functional processes of adults differs from that of adolescents, which may reflect the ongoing maturation of the limbic structures that regulate reactive aggression responses in adulthood.

Furthermore, the presence of *KRAS* gene as upstream regulator was also observed. Although *KRAS* is a proto-oncogene associated principally with cancer (Zhu et al., 2008), at the transcriptional level it acts as a short-term inductor to astrocytes in response to stimulus and as a sensor that adapts cells to metabolic needs and oxidative stress at the brain (Messina et al., 2017). The reactive aggression responses of the bulls during the *corrida* may trigger the mechanisms implicated in oxidative stress states, which have been observed to increase with age due to a disruption in the oxidant and anti-oxidant balance (Sies, 1991). Besides, in neuropsychiatric disorders such as bipolar disorder, elevated markers in blood of oxidative stress levels has been reported among adults, relating oxidative stress as mediator on neuropathological processes of adulthood (Andreazza et al., 2008).

Finally, it is worth also highlighting two genes implicated in functional development functions: the *NPAS4* gene, which participates in the structural and functional plasticity of neurons acting as regulator of the excitatory-inhibitory neural balance (Jaehne et al., 2015); and the *HTR2C* gene, modulator of the neural activity by functioning as down-stream effector of insulin sensitivity and glucose homeostasis (Stam et al., 1994). Both genes have also been candidates in studies of human pathophysiology in mental disorders, including schizophrenia, bipolar and cognitive disorders (Iwamoto et al., 2009), and are up (*NPAS4*) and down-regulated (*HTR2C*) in the adult group of bulls (Table 2).

Interestingly, whereas Agoglia et al. (2017) reported enhanced protein alterations in networks that regulate neuronal signaling, anxiety-related behavior and neurological disease in the adolescent PFC when compared to adult mice, our gene expression results found similar associations but only in adults. This different outcome may be due to differences in the age-stage comparison between studies. While in mice the biological transition of adolescence to adulthood is well studied, defining maturation stages in cattle has proved more complex, as its lifespan depends on the production purpose of each breed, among other factors (Essl, 1998). Given that the average life of a Lidia sire is approximately 15 years, and the average generation sire interval pathway (sire-offspring) is 7,5 years (Fernández-Salcedo, 1990), perhaps here the comparison between early and middle age adulthood would be a more correct approximation.

The present findings suggest that the cortical gene expression in 4-year-old Lidia cattle is characterized by an enhancement of molecular mechanisms involved in the increase of aggressive behaviors and neurophysiological disorders. However, despite our results suggesting cortical enhanced genetic expression associated with reactive aggression in adults, this may not be representative of other brain regions within the limbic system reported also as important actors in this process. Future experiments to evaluate the role of these other brain regions, particularly in the context of aggressive responses, will shed further light on this age transition period.

## Acknowledgements

The authors are grateful to the enterprise of Las Ventas, bullring “*Plaza 1*” and its veterinarian’s team for providing post-mortem Lidia breed brain tissue samples for this study.

